# *Pneumocystis* phosphoglucomutase links glucose-phosphate metabolism to β-1,3-glucan synthesis and cell-wall integrity

**DOI:** 10.64898/2026.11.04.764224

**Authors:** Theodore J. Kottom, Jenna L. McLean, Andrew H. Limper

## Abstract

The fungal cell wall is a dynamic, carbohydrate-rich structure whose composition is closely coordinated with cellular metabolism. Phosphoglucomutase (PGM) interconverts glucose-1- phosphate and glucose-6-phosphate, linking carbon metabolism with UDP-glucose-dependent cell-wall biosynthesis. We previously identified functional Pgm2 homologs from *Pneumocystis jirovecii* and *Pneumocystis murina*. Here, using a *Saccharomyces cerevisiae pgm2*Δ complementation model, we examined the relationship between Pgm2 activity, β-1,3-glucan, and cell-wall integrity. Pgm2 deficiency reduced β-1,3-glucan content and altered zymolyase sensitivity without major changes in cell-wall thickness. Bulk RNA sequencing revealed that the *pgm2*Δ mutant exhibited coordinated metabolic and cell-wall transcriptional responses, including increased *SLT2/MPK1*, *RLM1*, *FKS2/GSC2*, *GFA1*, *CHS1*, *CRH1*, *SED1*, and *CWP1* expression and decreased *FKS1*, *GAS1*, *GAS3*, and *CHS2* expression. Expression of either *Pjpgm*2 or *Pmpgm2* broadly shifted these responses toward wild-type levels. Selected transcriptional changes were further evaluated by qRT-PCR, which supported the major stress and cell-wall responses identified by RNA sequencing. *FKS2/GSC2* showed the same directional pattern, although differences were not statistically significant by qRT-PCR. Despite this compensatory response, Pgm2-deficient cells were markedly more susceptible to sublethal caspofungin, whereas either *Pneumocystis* Pgm2 homolog restored tolerance toward wild-type levels. These findings link Pgm2-dependent glucose-phosphate metabolism with β-1,3-glucan levels and cell-wall function. Given the importance of β-1,3-glucan to the *Pneumocystis* cyst wall and its limited cell-wall biosynthetic pathways, these findings further support Pgm2 as an important component of *Pneumocystis* cell-wall biology and a potential antifungal target.

**HIGHLIGHTS:**

- *Pneumocystis* Pgm2 restores β-1,3-glucan cell wall phenotypes in Pgm2-defecient yeast.
- Pgm2 loss remodels the fungal cell-wall integrity transcriptional program.
- Pgm2 deficiency induces *FKS2* and sensitizes yeast to caspofungin, a phenotype reversed by *Pneumocystis* Pgm2 homologs.

## INTRODUCTION

Pneumocystis jirovecii pneumonia (PJP) remains an important cause of morbidity and mortality among immunocompromised individuals (1, 2). A defining feature of the *Pneumocystis* ascus cell wall is its abundance of β-glucans, including β-1,3-glucan, which contribute to wall structure and host inflammatory recognition (3–6). Because β-glucan synthesis requires UDP-glucose (6), pathways that regulate glucose-phosphate metabolism may directly influence the ability of fungal cells to construct and remodel the cell wall.

Phosphoglucomutase (PGM) catalyzes the reversible interconversion of glucose-1-phosphate and glucose-6-phosphate (7). In *Saccharomyces cerevisiae*, *PGM2* encodes the major phosphoglucomutase activity, whereas *PGM1* provides a smaller residual contribution (8, 9).

We previously demonstrated that *P. jirovecii* and *P. murina* each encode a functional Pgm2 homolog and that *Pjpgm2* and *Pmpgm2* complement *S. cerevisiae pgm2*Δ phenotypes and restore Pgm2 activity (10, 11). Those findings genetically and biochemically validated *Pneumocystis* Pgm2 and suggested that the enzyme could represent a source of antifungal vulnerability. However, whether disruption of Pgm2 activity alters fungal cell-wall composition, cell-wall stress signaling, or susceptibility to agents targeting β-1,3-glucan synthesis remained unknown.

Fungal cell walls continuously remodel in response to metabolic and environmental stress. The cell-wall integrity network coordinates glucan and chitin synthesis, wall remodeling, and stress adaptation through pathways that include Rho1, Slt2/Mpk1, Rlm1, and Fks glucan synthases (12, 13). Because β-1,3-glucan synthesis depends on glucose-derived precursors (14), we reasoned that perturbing Pgm2 activity could alter cell wall composition while simultaneously triggering compensatory cell-wall signaling. Caspofungin blocks the production of β-1,3-glucan (15), making it a useful tool for studying how fungal metabolism affects cell wall maintenance.

Here, we integrate cell-wall ultrastructure, β-1,3-glucan measurements, zymolyase sensitivity, bulk RNA sequencing, confirmatory quantitative PCR, and caspofungin susceptibility to define how Pgm2 activity shapes fungal cell-wall composition and function. The data support a model in which loss of Pgm2 produces a metabolic defect that results in altered cell surface, activates compensatory cell-wall remodeling programs, and leaves the cell wall functionally vulnerable despite increased *FKS2* expression. Functional rescue by *P. jirovecii* and *P. murina pgm2* connects this cell-surface phenotype directly to *Pneumocystis* phosphoglucomutase activity.

## METHODS

### Yeast strains and culture conditions

The *Saccharomyces cerevisiae* strains used in this study were derived from the BY4742 background. The *pgm2*Δ strain was obtained from the *S. cerevisiae* deletion collection (Horizon Discovery) (16, 17). Wildtype (WT) and *pgm2*Δ strains transformed with pYES2.1/V5-His-TOPO/*lacZ* (Invitrogen) were used as vector controls. For complementation studies, the *Pneumocystis jirovecii pgm2* (*Pjpgm2*) and *Pneumocystis murina Pgm2* (*Pmpgm2*) genes were cloned into the pYES2.1/V5-His-TOPO expression vector and expressed in the *pgm2*Δ background. These strains and complementation constructs were previously generated and characterized in our earlier studies (10, 11). Yeast cultures were initially grown using raffinose as the carbon source and subsequently transferred to galactose-containing medium to induce expression from the *GAL1* promoter. The four experimental groups used throughout the study were wild-type (WT) + vector, *pgm2*Δ + vector, *pgm2*Δ + *Pjpgm2*, and *pgm2*Δ + *Pmpgm2*.

### Transmission electron microscopy and cell-wall analysis

Yeast cultures were harvested by centrifugation (8,000 rpm, 5 min) and fixed for 24 h at 4°C in 4% paraformaldehyde and 2% glutaraldehyde in 0.1 M phosphate buffer (pH 7.0). Fixed cells were embedded in 1% low-melting-point agar, washed in 0.15 M cacodylate buffer, post-fixed with 2% osmium tetroxide, stained with 2% aqueous uranyl acetate, dehydrated through a graded ethanol series, and embedded in Spurr resin. Ultrathin sections (100 nm) were stained with lead citrate and examined using an HT7800 120-kV transmission electron microscope (Hitachi High-Tech). Cell-wall thickness was measured from TEM images obtained at 40,000X magnification. Only cells with clearly visible wall boundaries and suitable section orientation were included. Damaged, partially cut, poorly defined, or angled wall regions were excluded. Wall thickness was measured at multiple locations around each suitable cell, and these measurements were combined to obtain a representative thickness for each cell. All cell measurements were obtained from a single TEM experiment and therefore represent individual cells rather than independent experiments.

### β-1,3-glucan quantification

Cellular β-1,3-glucan content was measured using an aniline blue fluorescence assay adapted from previously described methods (18, 19). Equivalent numbers of WT + vector, *pgm2*Δ + vector, *pgm2*Δ + *Pjpgm2*, and *pgm2*Δ + *Pmpgm2* cells were harvested and washed twice with TE buffer (10 mM Tris-HCl, 1 mM EDTA, pH 8.0). Cells were resuspended in 500 μL TE buffer, mixed with 100 μL of 6 M NaOH, and incubated at 80°C for 30 min to solubilize cell-wall glucan.

Aniline blue reagent (2.1 mL; 0.03% aniline blue, 0.18 M HCl, and 0.49 M glycine-NaOH, pH 9.5) was then added, and samples were incubated at 50°C for 30 min followed by 30 min at room temperature. Fluorescence was measured using a fluorescence plate reader at approximately 400 nm excitation and 460 nm emission. Background fluorescence was subtracted, and β-1,3-glucan levels were normalized to the WT control and expressed as percent of WT.

### Zymolyase cell-wall susceptibility assay

Cell-wall susceptibility to enzymatic digestion was assessed using a zymolyase sensitivity assay (20–22). WT + vector, *pgm2*Δ + vector, *pgm2*Δ + *Pjpgm2*, and *pgm2*Δ + *Pmpgm2* cultures were harvested and washed, and cell suspensions were adjusted to the same starting optical density (OD600). Reactions were prepared in 1X storage buffer supplied with the enzyme and treated with 0.5 U of Zymolyase (Zymo Research) per reaction. OD600 was measured immediately before treatment and at the indicated time points during digestion. Cell-wall digestion was expressed as the percentage of the initial OD600 remaining at each time point, with the starting OD600 defined as 100%. A greater decrease in OD600 indicated increased susceptibility to cell-wall digestion, whereas retention of OD600 indicated greater resistance to zymolyase.

### Bulk RNA sequencing and transcriptomic analysis

Total RNA was isolated from four independent clones each of WT + vector, *pgm2*Δ + vector, *pgm2*Δ + *Pjpgm2*, and *pgm2*Δ + *Pmpgm2* yeast (16 samples total). Cells were mechanically disrupted using a QIAGEN TissueLyser and 7 mm stainless steel beads, and RNA was purified using the RNeasy Plus Mini Kit (QIAGEN) according to the manufacturer’s protocol. Genomic DNA was removed during RNA purification using the genomic DNA removal procedure included with the kit, followed by on-column DNase digestion as recommended by the manufacturer. Purified RNA was submitted to Plasmidsaurus for DGE RNA-Seq. Sequencing generated approximately 18–21 million reads per sample. Reads were quality filtered using fastp (v0.24.0), and read quality was assessed using FastQC (v0.12.1). Filtered reads were aligned to the *Saccharomyces cerevisiae* reference genome (R64-1-1_114) using STAR (v2.7.11b), followed by coordinate sorting with samtools (v1.21). PCR duplicates were removed by UMI-based deduplication using UMIcollapse (v1.1.0). Gene-level expression was quantified using feature Counts from the Subread package (v2.1.1) with strand-specific counting and reported as counts per million (CPM). Sample quality and reproducibility were assessed using read-mapping metrics, sample-to-sample correlation, and principal component analysis. Differential expression was evaluated using the Plasmidsaurus edge python quasi-likelihood workflow, including expression filtering, normalization, dispersion estimation, model fitting, and quasi-likelihood testing. For visualization of selected genes, mean CPM values were calculated from the four independent clones within each group. Heat-map colors represent row-wise Z-scores calculated from the group mean CPM values, while the corresponding mean CPM values are displayed within each cell. Based on the bulk RNA-seq findings, selected genes representing stress and carbon responses, cell-wall integrity signaling, β-1,3-glucan synthesis, cell-wall remodeling, and structural cell-wall proteins were chosen for focused expression analysis. Expression of the *Pjpgm2* and *Pmpgm2* transgenes was also quantified to verify expression in the corresponding *pgm2*Δ complementation strains. Representative genes from the major transcriptional response pathways identified by RNA sequencing were subsequently evaluated by qRT-PCR to independently assess these expression patterns. Raw and processed RNA-sequencing data generated in this study have been deposited in the NCBI Gene Expression Omnibus (GEO) and Sequence Read Archive (SRA) under accession number [GSE/PRJNA accession number].

### Quantitative RT-PCR

Selected genes identified by bulk RNA sequencing were independently evaluated by quantitative reverse-transcription PCR (qRT-PCR). Genes were selected to represent stress and carbon responses (*ADY2, HSP12*, *HXT2*, *HXT5*, and *NQM1*), cell-wall integrity signaling (*RHO1*, *RLM1, SLT2/MPK1*), β-1,3-glucan synthesis (*FKS1* and *FKS2/GSC2*), and cell-wall remodeling/structural (*CRH1*, *PIR3*, and *GAS1*) (23–25). Total RNA was reverse transcribed to cDNA using the iScript Advanced cDNA Synthesis Kit for RT-qPCR (Bio-Rad) according to the manufacturer’s instructions. The equivalent of 50 ng of cDNA was used as template for each qPCR reaction. Reactions were performed in duplicate in a final volume of 12.5 µL/well using SsoAdvanced SYBR Green chemistry (Bio-Rad), with gene-specific forward and reverse primers at a final concentration of 500 nM each. Primer sequences are provided in Supplemental Table S1. Amplification was performed using a Bio-Rad CFX96 Real-Time PCR Detection System according to the manufacturer’s recommended thermocycling conditions, followed by melt-curve analysis to assess amplification specificity. Gene expression was normalized to *ACT1*, and relative expression was calculated using the comparative Ct (2^-ΔΔCt) method (26, 27). Technical duplicates were averaged for each sample before analysis.

### Caspofungin susceptibility assay

Caspofungin susceptibility was evaluated in WT + vector, *pgm2*Δ + vector, *pgm2*Δ + *Pjpgm2*, and *pgm2*Δ + *Pmgm2* yeast (28, 29). Cultures were initially grown in selective medium containing raffinose and subsequently induced with galactose under identical conditions prior to drug exposure. Following induction, cultures were adjusted to the same starting OD600 and maintained in galactose-containing medium throughout the assay. Caspofungin acetate (Cayman Chemical) was prepared in sterile PBS and tested over a concentration range of 0–1 µg/mL. Reactions were performed in 96-well plates at a final volume of 200 µL per well, with each strain and drug concentration tested in triplicate. Vehicle-control wells received an equivalent volume of PBS. OD600 was monitored during growth with shaking between measurements. Initial dose-response experiments were used to identify a low caspofungin concentration that permitted WT growth while revealing differential susceptibility among strains. For the final analysis, growth at 12 h in the presence of 0.063 µg/mL caspofungin was compared with the corresponding vehicle-treated culture for each strain. Growth in the vehicle control was defined as 100%, and caspofungin-treated growth was expressed relative to the matched untreated control. Four independent experiments were analyzed, with technical replicates averaged within each experiment before statistical analysis.

### Statistical Analysis

Data were analyzed by one-way ANOVA to determine differences across experimental conditions, with two-tailed Student’s t-tests used for appropriate pairwise comparisons. Analyses were performed in GraphPad Prism 11.1.0 (GraphPad Software, San Diego, CA), and P values below 0.05 were considered statistically significant.

## RESULTS

### *pgm2*Δ deletion and *Pneumocystis pgm2* complementation do not substantially alter overall cell-wall thickness

Transmission electron microscopy was used to examine cell-wall ultrastructure in WT + vector, *pgm2*Δ + vector, and *pgm2*Δ strains complemented with either *Pjpgm2* or *Pmpgm2* (Fig. 1A-D). All four strains displayed a clearly defined cell wall, although some variability in wall appearance was observed among individual cells. Quantitative analysis of the 40,000X images showed similar overall wall thickness among the four strains (Fig. 2). Mean ± SD thickness was 91.4 ± 20.2 nm for WT + vector (n = 6 cells), 87.7 ± 30.7 nm for *pgm2*Δ + vector (n = 6), 100.5 ± 20.2 nm for *Pjpgm2*-complemented cells (n = 9), and 90.7 ± 17.8 nm for *Pmpgm2*-complemented cells (n = 8). Thus, loss of Pgm2 or complementation with either *Pneumocystis* Pgm2 homolog did not produce an obvious change in overall cell-wall thickness. We next assessed β-1,3-glucan abundance and functional cell-wall integrity.

**Figure 1.**
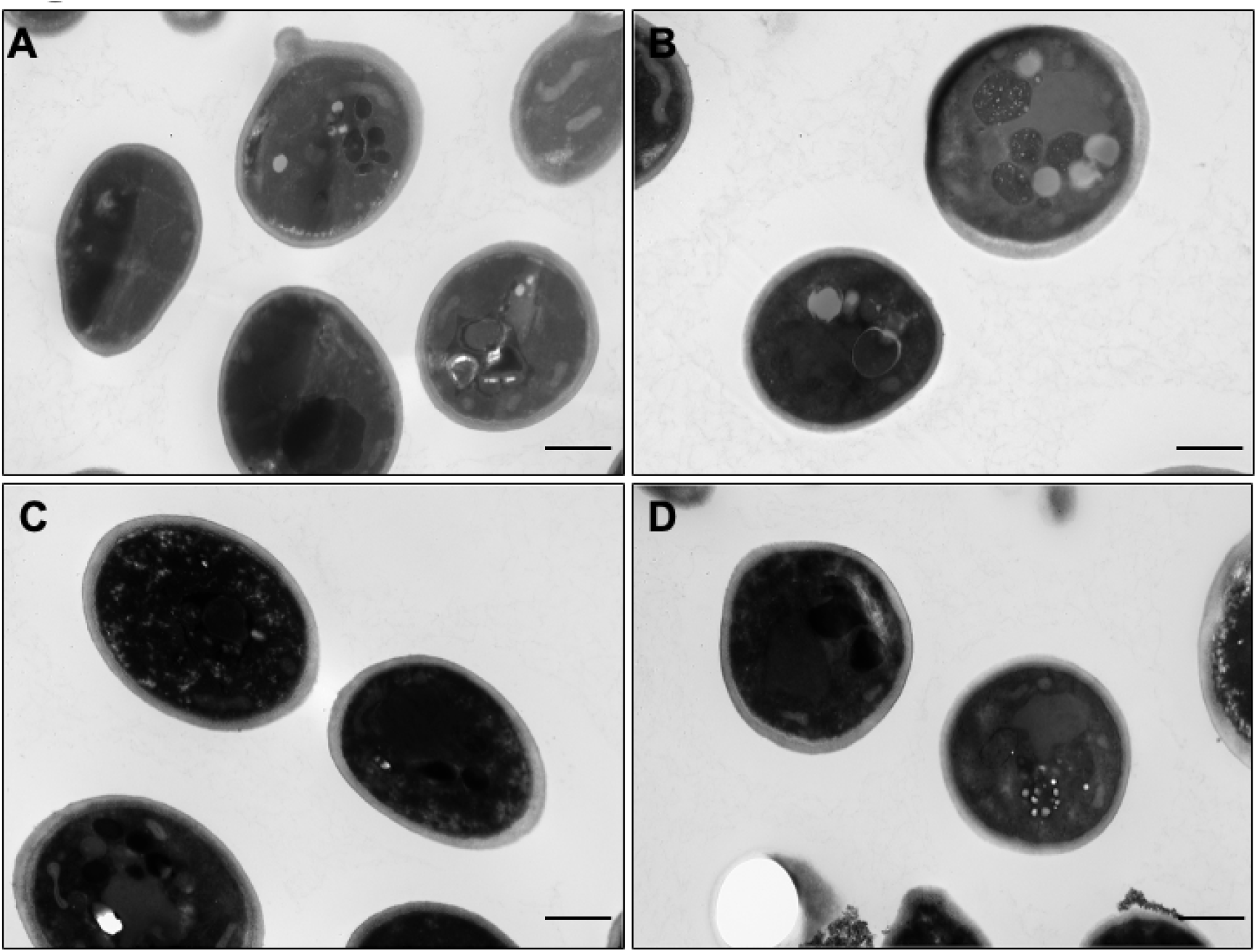
Transmission electron microscopy of WT, Δ*pgm2*, and complemented strains. Representative transmission electron microscopy (TEM) images of **(A)** WT + vector, **(B)** Δ*pgm2* + vector, **(C)** Δ*pgm2* + *Pjpgm2*, and (D) Δ*pgm2* + *Pmpgm2*. Images were acquired at 40,000X magnification. Scale bars, 800 nm.

**Figure 2.**
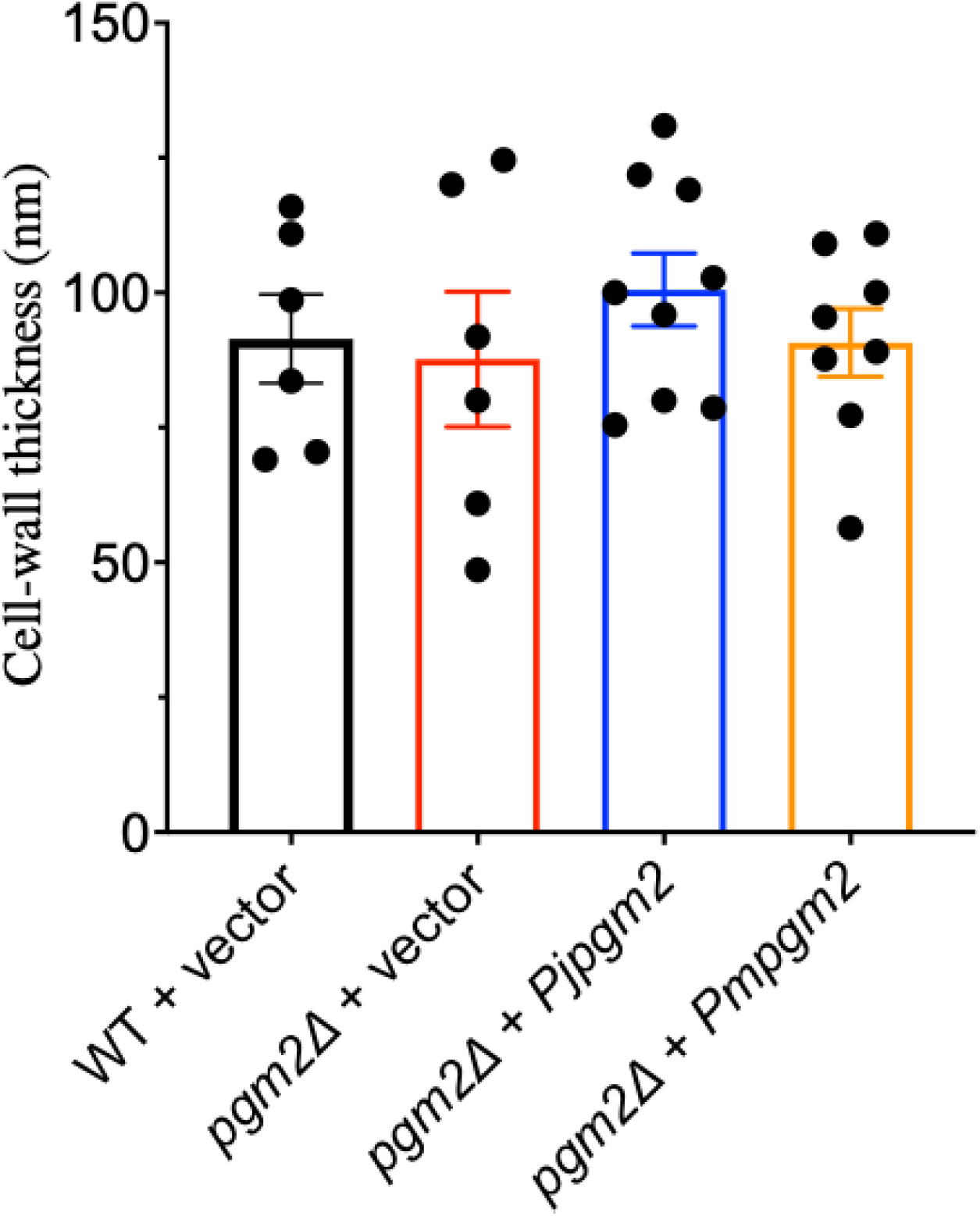
Cell wall thickness of WT, Δ *pgm2*, and complemented strains. Cell wall thickness was measured from transmission electron microscopy (TEM) images of WT + vector, Δ*pgm2* + vector, Δ*pgm2* + *Pjpgm2*, and Δ*pgm2* + *Pmpgm2*. Bars represent the mean ± SD, with individual biological replicates shown as points. No statistically significant differences in cell wall thickness were observed among the strains.

### Pgm2 deficiency reduces β-1,3-glucan content despite preservation of overall cell-wall architecture

TEM analysis did not reveal major changes in cell-wall ultrastructure or thickness. We next examined cell-wall composition to determine whether Pgm2 deficiency affected other features of cell-wall architecture. β-1,3-glucan content was quantified by aniline blue fluorescence and normalized to wild-type levels. The *pgm2*Δ + vector strain showed a significant reduction in β-1,3-glucan, to approximately 84% of WT + vector levels (Fig. 3; ****P < 0.0001). Thus, despite the absence of an obvious change in overall wall thickness by TEM, loss of Pgm2 reduced a major structural component of the fungal cell wall. Expression of either *Pjpgm2* or *Pmpgm2* in the *pgm2*Δ background restored β-1,3-glucan content to approximately WT + vector levels. Both complemented strains contained significantly more β-1,3-glucan than the *pgm2*Δ + vector strain (****P < 0.0001) and were comparable to WT + vector. These findings indicate that Pgm2 activity is required to maintain normal β-1,3-glucan levels and demonstrate that both *Pneumocystis* Pgm2 homologs can functionally restore this phenotype.

**Figure 3.**
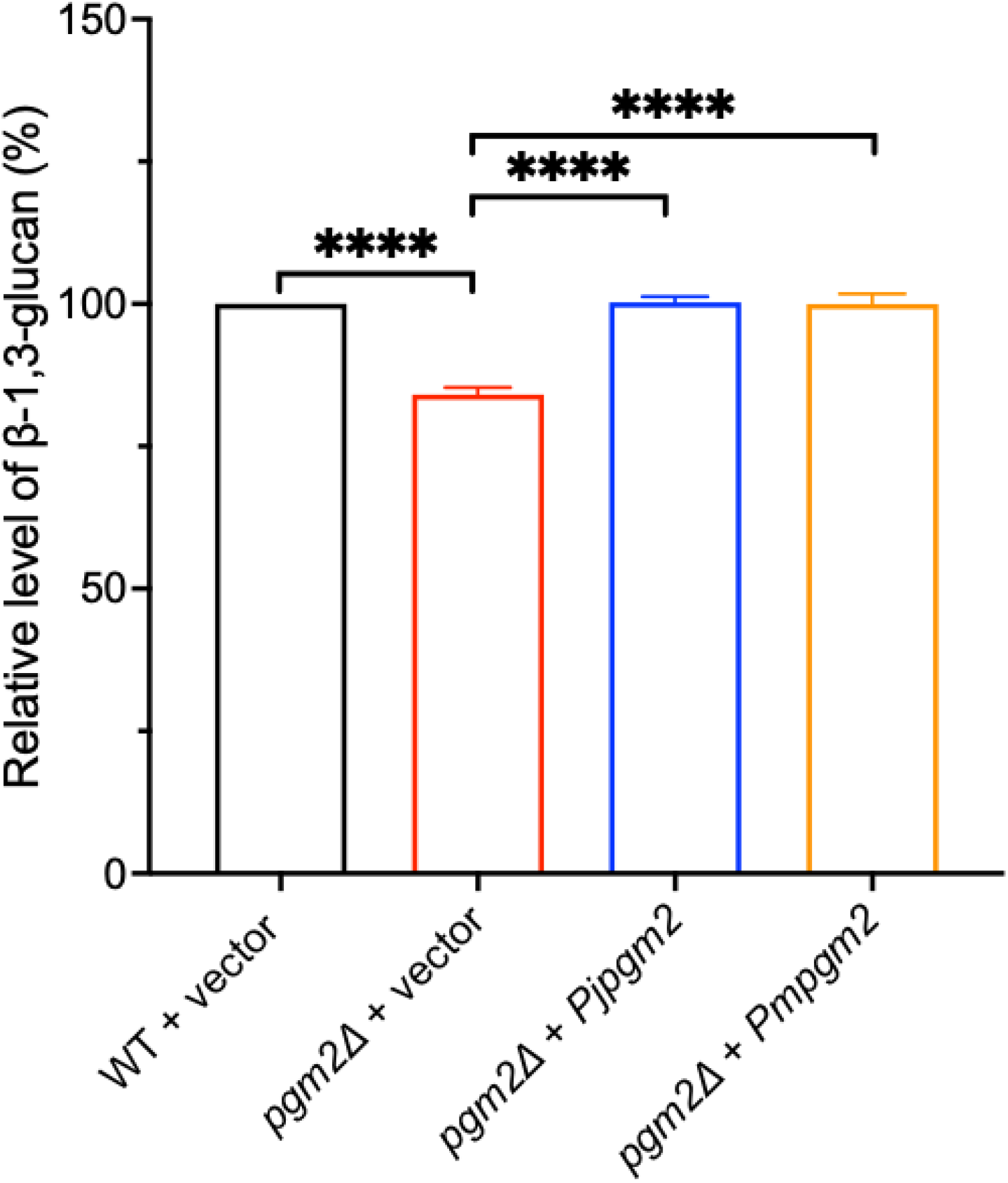
Relative β-1,3-glucan levels in WT, Δ*pgm2*, and complemented strains. Relative β-1,3-glucan levels were determined using an aniline blue fluorescence assay in WT + vector, Δ*pgm2* + vector, Δ*pgm2* + *Pjpgm2*, and Δ*pgm2* + *Pmpgm2*. Values were normalized to the WT + vector control within each experiment, which was set to 100%. Data were obtained from three independent experiments, with three technical replicates per experiment, and are presented as the mean ± SD. Statistical significance was assessed by one-way ANOVA followed by t-tests. ****P < 0.0001.

### Pgm2 deficiency compromises functional cell-wall integrity

To determine whether the reduction in β-1,3-glucan associated with Pgm2 deficiency resulted in a functional alteration of the cell wall, sensitivity to enzymatic cell-wall disruption was assessed using zymolyase. Previous studies have shown that β-glucan is accessible to zymolyase in WT cells but less accessible in several yeast cell-wall mutants (30, 31). Thus, susceptibility to zymolyase provides a functional measure of β-glucan accessibility within the cell-wall architecture. Cell-wall digestion was monitored over time by measuring the decrease in optical density (OD600). WT + vector cells exhibited a progressive decrease in OD600 during zymolyase treatment, consistent with enzymatic degradation of the cell wall (Fig. 4). In contrast, *pgm2*Δ cells containing the vector control displayed markedly reduced zymolyase-mediated digestion, retaining a substantially greater proportion of their initial optical density throughout the assay. Thus, despite having reduce β-1,3-glucan content, Pgm2-deficent cells were more resistant to zymolyase-mediated cell-wall disruption. Expression of either *Pjpgm2* or *Pmpgm2* reversed this phenotype. These findings suggest that the *Pneumocystis* Pgm2 homologs restore not only β-1,3-glucan levels but also aspects of its function in fungal cell architecture. Despite reduced β-1,3-glucan, *pgm2*-deficient cells were more resistant to zymolyase, suggesting compensatory remodeling rather than loss of the carbohydrate. We therefore examined the transcriptional response to Pgm2 absence to identify pathways associated with cell-wall modeling and if the *Pneumocystis* Pgm2 homologs could restore this defect.

**Figure 4.**
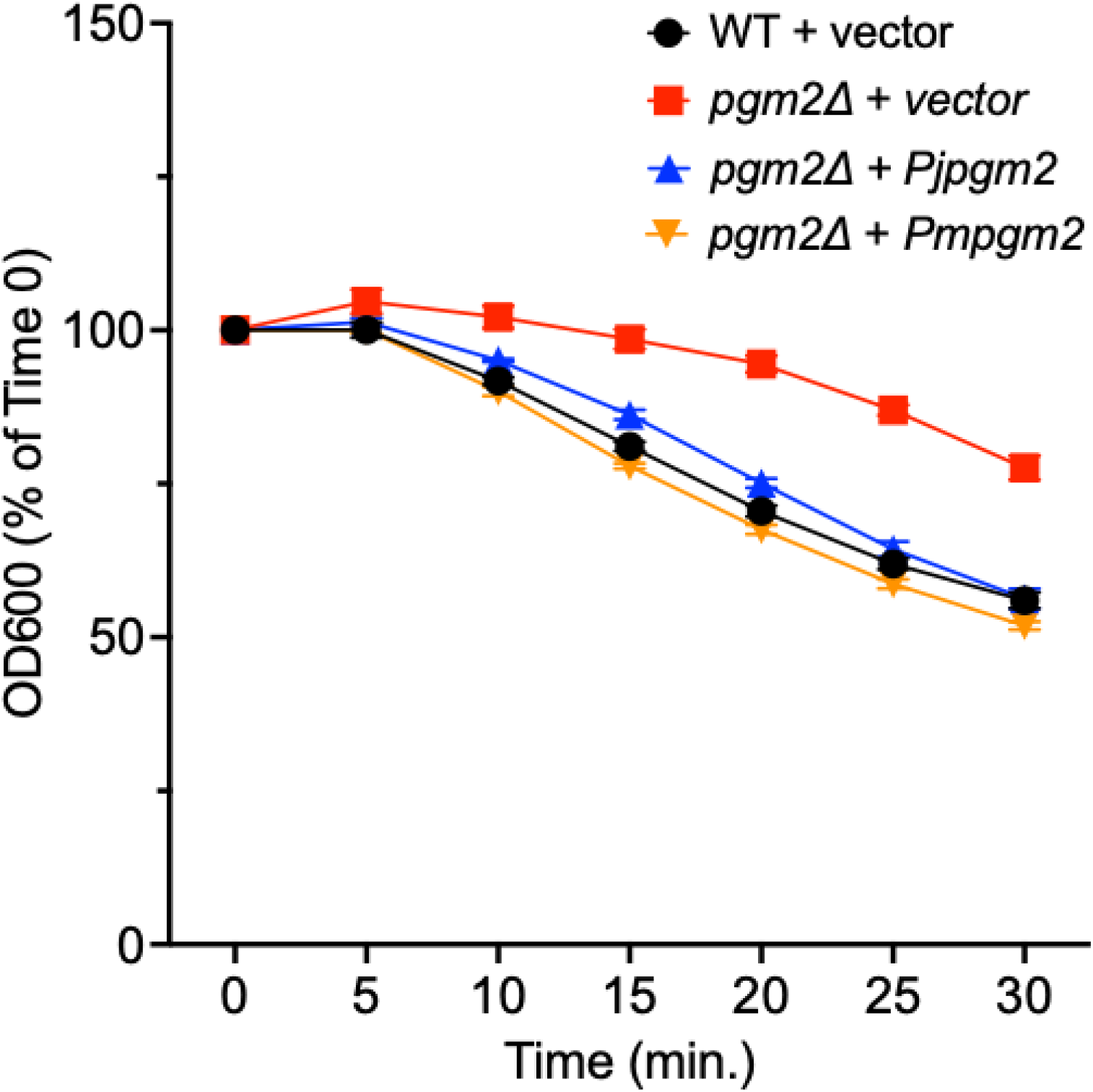
Zymolyase susceptibility of WT, Δ*pgm2*, and complemented strains. Susceptibility to zymolyase-mediated cell wall digestion was assessed in WT + vector, Δ*pgm2* + vector, Δ*pgm2* + *Pjpgm2*, and Δ*pgm2* + *Pmpgm2* by monitoring the decrease in optical density over time. Optical density at each time point was expressed relative to the initial value at 0 min (100%). For each independent experiment, values were averaged from triplicate technical wells. Data represent the mean ± SD from two independent experiments.

### Pgm2 deficiency induces coordinated metabolic and cell-wall transcriptional responses that are reversed by *Pneumocystis pgm2* complementation

Bulk RNA sequencing was performed on four independent clones each of WT + vector, *pgm2*Δ + vector, *pgm2*Δ + *Pjpgm2*, and *pgm2*Δ + *Pmpgm2* yeast. Sample-to-sample correlation analysis showed highly similar transcriptional profiles among the four independent clones within each experimental group, with Pearson correlation coefficients of approximately 0.994–0.997.

These strong within-group correlations indicate high reproducibility across independent clones and support the consistency of the RNA-seq dataset. In contrast, correlations between experimental groups were lower, with the greatest difference observed between WT + vector and *pgm2*Δ + vector samples (Fig. 5). The *Pjpgm2*- and *Pmpgm2*-complemented strains exhibited transcriptional profiles that shifted away from the *pgm2*Δ + vector profile and toward WT + vector, consistent with broad restoration of gene expression following complementation. Transgene analysis confirmed robust expression of the corresponding *Pneumocystis pgm2* homolog in each complemented strain. Mean *Pjpgm2* expression was approximately 24,698 CPM in the *Pjpgm2*-complemented clones, while mean *Pmpgm2* expression was approximately 13,966 CPM in the *Pmpgm2*-complemented clones, with only minimal background signal detected in the other groups (Fig. 6). Pathway-focused analysis revealed coordinated transcriptional changes involving carbon metabolism and cellular stress, cell-wall integrity signaling, β-1,3-glucan synthesis, chitin synthesis and precursor pathways, cell-wall remodeling, and cell-wall structural proteins (Fig. 6). The complete gene-level expression dataset across all experimental groups is available in the deposited GEO dataset. The stress- and carbon-responsive genes *HSP12*, *HXT2*, *ADY2*, *HXT5*, and *NQM1* were strongly increased in *pgm2*Δ + vector cells compared with WT + vector cells, whereas expression in both complemented strains shifted toward WT + vector levels. Pgm2 deficiency was also associated with increased expression of genes involved in the cell-wall integrity response, including *RHO1*, *SLT2/MPK1*, and *RLM1*, together with the cell-wall remodeling genes *CRH1* and *PIR3*. *FKS2/GSC2*, encoding the stress-responsive β-1,3-glucan synthase subunit, was also strongly induced. In contrast, *GAS1*, which participates in β-1,3-glucan remodeling (30), was reduced following *pgm2* deletion. Expression of either *Pneumocystis pgm2* homolog broadly shifted these transcriptional changes toward the WT + vector state. Together, the sample-to-sample correlation and pathway-focused expression analyses indicate that loss of Pgm2 is associated with a coordinated transcriptional response involving metabolic stress and multiple aspects of cell-wall function. Expression of either *Pjpgm2* or *Pmpgm2* broadly shifted these responses toward the WT + vector pattern. Representative genes from these pathways were subsequently evaluated by qRT-PCR to independently assess the expression patterns identified by bulk RNA sequencing.

**Figure 5.**
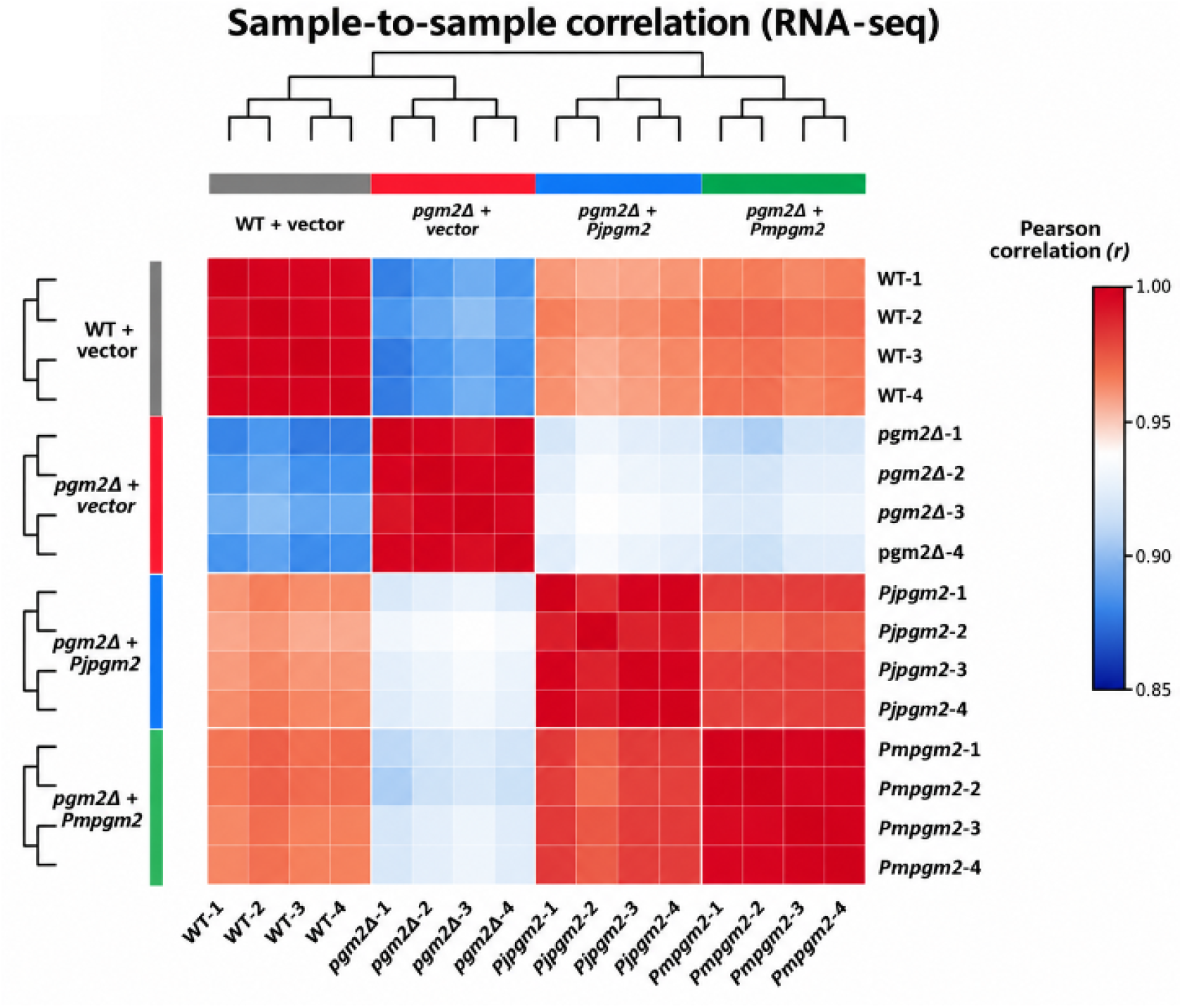
Sample-to-sample correlation analysis of RNA-seq datasets from WT, Δ*pgm2*, and complemented strains. Pairwise Pearson correlation coefficients (r) were calculated from RNA-seq expression profiles for WT + vector, Δ*pgm2* + vector, Δ*pgm2* + *Pjpgm2*, and Δ*pgm2* + *Pmpgm2*, with four biological replicates per strain. Correlation coefficients are represented by the color scale, with warmer colors indicating greater similarity between samples. Hierarchical clustering based on sample-to-sample correlations is shown by the dendrograms. Biological replicates clustered closely within each strain, while the Δ*pgm2* + vector samples formed a distinct cluster from WT + vector and the complemented strains.

**Figure 6.**
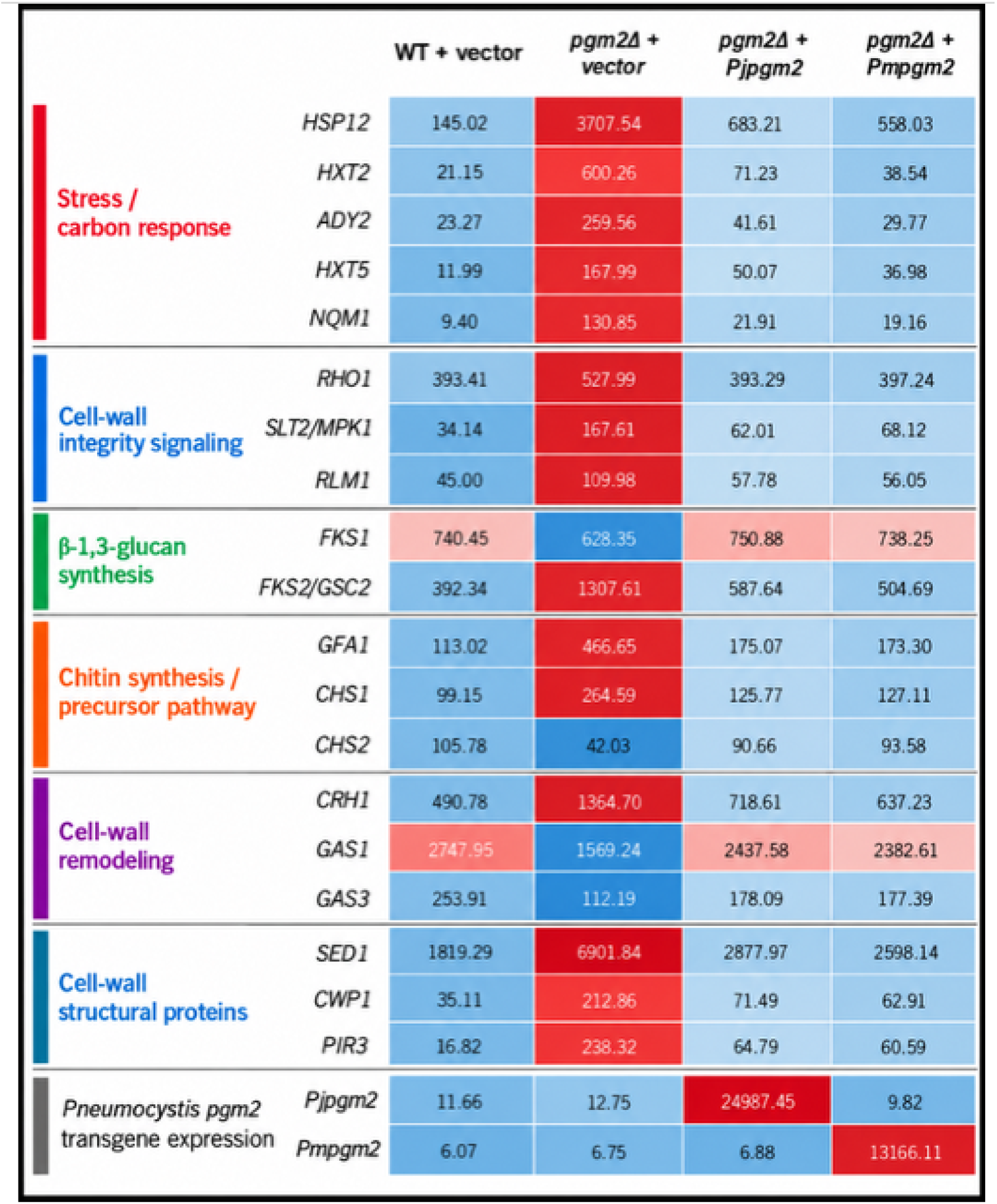
RNA-seq expression profiles of selected cell wall-associated and stress-response genes in WT + vector, Δ*pgm2* + vector, and complemented strains. Mean RNA-seq expression levels are shown for selected genes associated with stress/carbon response, cell-wall integrity signaling, β-1,3-glucan synthesis, and cell-wall remodeling in WT + vector, Δ*pgm2* + vector, Δ*pgm2* + *Pjpgm2*, and Δ*pgm2* + *Pmpgm2*. Values represent mean counts per million (CPM) across four independent clones for each strain. Expression of the *Pjpgm2* and *Pmpgm2* transgenes is shown separately at the bottom as mean CPM. Heatmap coloring represents relative expression across strains for each gene, with the numerical mean CPM displayed within each cell.

### qRT-PCR supports Pgm2-dependent stress and cell-wall transcriptional responses

To independently evaluate the transcriptional changes identified by bulk RNA sequencing, representative genes involved in stress and carbon metabolism, cell-wall integrity signaling, β-1,3-glucan synthesis, cell-wall remodeling, and cell-wall structure were examined by qRT-PCR (Fig. 7). Individual biological replicates and statistical comparisons are shown in Fig. 7A, while the corresponding mean relative-expression values are summarized in Fig. 7B, with row Z-scores used for heat-map coloration. Consistent with the RNA-seq findings, the stress-and carbon-responsive genes *HSP12*, *HXT2*, *ADY2*, *HXT5*, and *NQM1* were increased in *pgm2*Δ + vector cells compared with WT + vector cells. Expression of each gene shifted toward WT + vector levels following complementation with either *Pjpgm2* or *Pmpgm2*. Overall differences among the four groups were significant by one-way ANOVA for the stress-and carbon-responsive genes examined (Fig. 7A), supporting a transcriptional response associated with Pgm2 deficiency that was largely reversed by expression of either *Pneumocystis pgm2* homolog. Genes associated with cell-wall integrity signaling showed a similar overall pattern. *SLT2/MPK1* was strongly increased in *pgm2*Δ + vector cells and shifted toward WT + vector levels in both complemented strains. *RHO1* and *RLM1* showed more modest increases following *pgm2* deletion, with expression shifting toward WT + vector levels following complementation. Among genes associated with cell-wall remodeling and structure, *CRH1* and the structural cell-wall protein gene *PIR3* were increased in *pgm2*Δ + vector cells and shifted toward WT + vector levels following *Pjpgm2* or *Pmpgm2* complementation. In contrast, *GAS1* showed the opposite pattern, with reduced expression in *pgm2*Δ + vector cells and recovery toward WT + vector levels following complementation. Together, these findings support coordinated transcriptional changes in genes associated with cell-wall integrity signaling, remodeling, and structure following loss of Pgm2. The β-1,3-glucan synthase genes showed more modest changes by qRT-PCR. *FKS1* expression was lower in *pgm2*Δ + vector cells and remained lower in the complemented strains, consistent with the direction observed by bulk RNA sequencing; however, differences among groups did not reach statistical significance (Fig. 7B). *FKS2/GSC2* also showed a directional pattern consistent with the bulk RNA-seq data, increasing from a mean relative expression of 0.949 in WT + vector cells to 1.352 in *pgm2*Δ + vector cells and shifting toward WT + vector levels following *Pjpgm2* (1.091) or *Pmpgm2* (1.047) complementation. Although this pattern paralleled that observed by RNA sequencing, the overall difference among groups was not statistically significant by qRT-PCR (one-way ANOVA, P = 0.21). Thus, the qRT-PCR data did not establish significant differences in *FKS1* or *FKS2/GSC2* expression but showed directional patterns consistent with those observed by bulk RNA sequencing.

**Figure 7.**
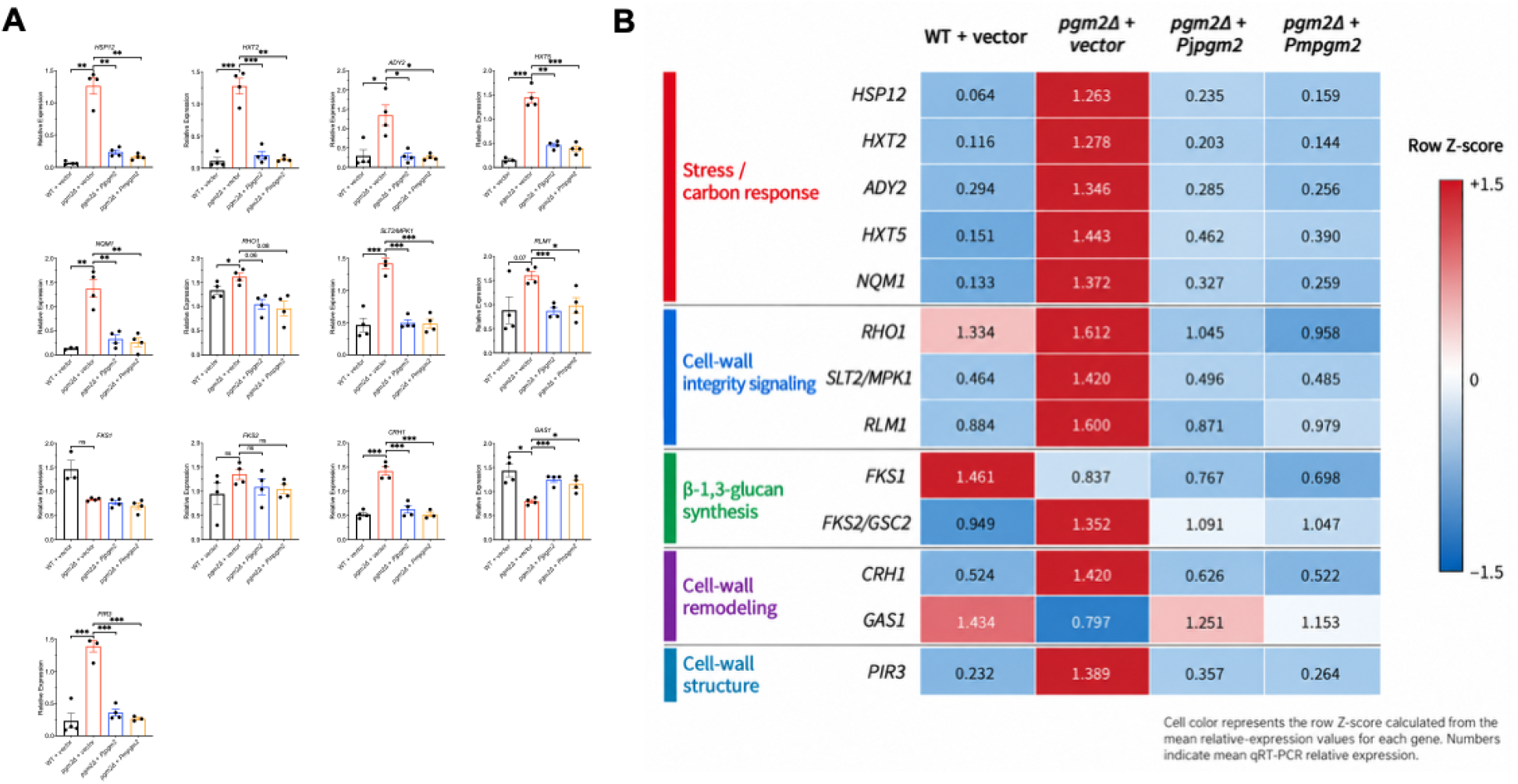
qRT-PCR analysis of stress- and cell wall-associated gene expression in WT + vector, *pgm2*Δ + vector, and complemented strains. **(A)** Relative expression of genes associated with stress and carbon metabolism, cell-wall integrity signaling, β-1,3-glucan synthesis, cell-wall remodeling, and cell-wall structure was measured by qRT-PCR in WT + vector, *pgm2*Δ + vector, *pgm2*Δ + *Pjpgm2*, and *pgm2*Δ + *Pmpgm2* strains. Data represent the mean ± SD from 3–4 independent experiments, with individual experimental values shown as points. Statistical significance is indicated above the corresponding comparisons. **(B)** Heat map summarizing the qRT-PCR expression data shown in panel A. Numerical values within each cell represent the mean relative expression for each gene and strain. Cell color represents the row Z-score calculated from the mean relative-expression values for each gene and was used only for heat-map visualization, with red indicating relatively higher expression and blue indicating relatively lower expression within each gene.

**Figure 8.**
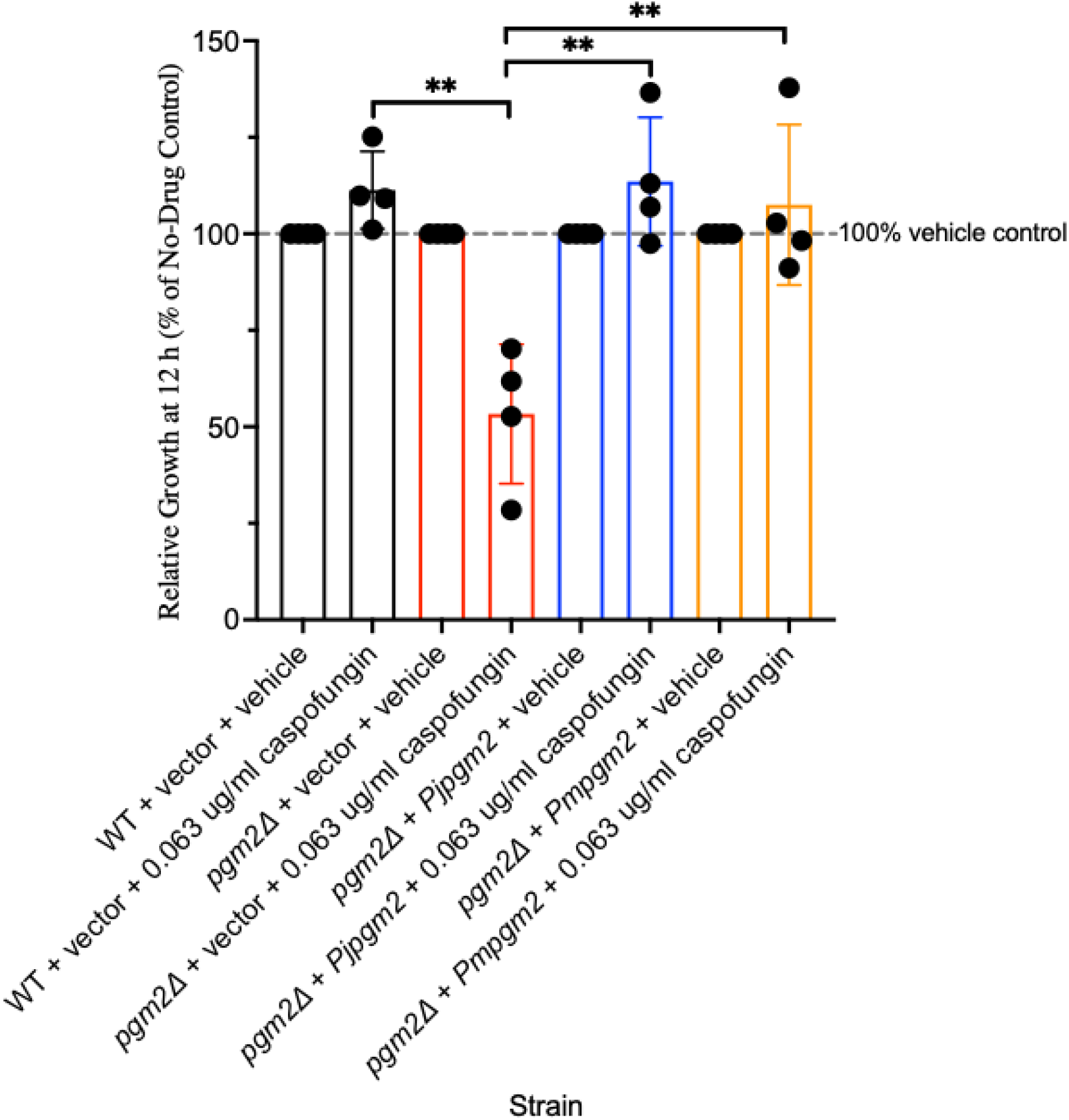
Caspofungin sensitivity of WT, Δ*pgm2*, and complemented strains. Caspofungin sensitivity was assessed in WT + vector, Δ*pgm2* + vector, Δ*pgm2* + *Pjpgm2*, and Δ*pgm2* + *Pmgm2* following 12 h of exposure of 0.063ug/mL caspofungin. Growth in the presence of caspofungin was expressed as a percentage of the corresponding no-drug control for each strain. Data represent the mean ± SD from four independent experiments, with each experiment performed in triplicate wells. Statistical significance is indicated above the corresponding comparisons. **P < 0.01.

Overall, qRT-PCR supported the major transcriptional patterns identified by bulk RNA sequencing, including stress and carbon responses and changes in genes associated with cell-wall integrity signaling, remodeling, and structure following Pgm2 deficiency. Expression of either *Pneumocystis* Pgm2 homolog broadly shifted many of these responses toward the WT state. Although the qRT-PCR analysis did not establish significant changes in *FKS1* or *FKS2/GSC2*, the directional increase in *FKS2/GSC2* following Pgm2 deficiency and its shift toward WT + vector levels following complementation paralleled the RNA-seq findings. Considered together with the reduction in β-1,3-glucan abundance and transcriptional changes in cell-wall integrity-associated genes, these findings suggested that Pgm2 deficiency may be accompanied by a compensatory response involving the β-1,3-glucan synthesis pathway. We therefore next examined the response of Pgm2-deficient cells to caspofungin, an echinocandin inhibitor of β-1,3-glucan synthase.

### Pgm2 deficiency increases susceptibility to β-1,3-glucan synthase inhibition

The increase in *FKS2/GSC2* expression observed in *pgm2*-deficient cells, together with reduced β-1,3-glucan abundance and activation of the cell-wall integrity response, suggested that loss of Pgm2 may increase dependence on compensatory β-1,3-glucan synthesis. We therefore examined the response of the four strains to caspofungin, an echinocandin inhibitor of β-1,3-glucan synthase. Following an initial dose-response analysis, growth was compared at 12 h using 0.063 µg/mL caspofungin, a concentration that permitted continued WT + vector growth while revealing a marked phenotype in *pgm2*-deficient cells. Growth was normalized to the corresponding vehicle-treated culture for each strain. WT + vector cells retained approximately 111% of vehicle-control growth, whereas growth of the *pgm2*Δ + vector strain was reduced to approximately 53%. In contrast, complementation with either *Pjpgm2* or *Pmpgm2* restored growth in the presence of caspofungin to approximately 114% and 107% of the corresponding vehicle controls, respectively. Thus, Pgm2 deficiency produced a marked increase in caspofungin susceptibility that was reversed by expression of either *Pneumocystis* Pgm2 homolog. Together with the reduced β-1,3-glucan content and the transcriptional activation of cell-wall integrity and *FKS2/GSC2*-associated responses, these findings support a model in which loss of Pgm2 compromises β-1,3-glucan homeostasis and increases reliance on the Fks-dependent compensatory response.

## DISCUSSION

Our previous studies established that *Pneumocystis jirovecii* and *Pneumocystis murina* encode functional phosphoglucomutases capable of complementing Pgm2 deficiency in *Saccharomyces cerevisiae* (10, 11). We further demonstrated that inhibition of PGM activity reduces *Pneumocystis* viability and that the fungal-selective PGM inhibitor ISFP10 inhibits *Pneumocystis* PGM activity (10, 30, 31). The present study extends these findings by demonstrating that PGM activity is functionally linked to β-1,3-glucan homeostasis, cell-wall stress signaling, and susceptibility to β-1,3-glucan synthase inhibition. Together, these observations suggest that PGM is not simply a metabolic enzyme required for fungal growth, but also as an upstream determinant of fungal cell-wall integrity.

Loss of Pgm2 produced a substantial reduction in β-1,3-glucan despite relatively preserved overall cell-wall ultrastructure and thickness. The *pgm2*Δ strain was also more resistant to zymolyase digestion, suggesting that Pgm2 deficiency alters wall composition, organization, or enzyme accessibility rather than simply decreasing total wall material. Both biochemical phenotypes were restored toward WT + vector by expression of either *Pjpgm2* or *Pmpgm2*. These findings are consistent with the metabolic position of PGM between glucose-6-phosphate and glucose-1-phosphate, the latter supporting UDP-glucose production and thereby providing substrate for β-glucan synthesis (32, 33). Our previous characterization of *Pneumocystis* Pgm2 proposed this connection between PGM-dependent carbohydrate metabolism, UDP-glucose availability, and the link to the glucan-rich cyst wall (10, 11).

The importance of this metabolic connection is particularly relevant to *Pneumocystis*. β-1,3-glucan is a major structural component of the thick-walled cyst form but is essentially absent from trophic forms (4, 34). Earlier studies identified and functionally characterized the *P. carinii gsc1* gene encoding the catalytic β-1,3-glucan synthase subunit and demonstrated strong life-cycle regulation, with *gsc1* expression and β-1,3-glucan synthesis concentrated in the cyst form (6). β-1,3-glucan synthase inhibitors likewise markedly reduce cyst numbers in experimental infection (35–37). Thus, β-glucan synthesis is linked not only to wall structure but also to *Pneumocystis* life-cycle biology and transmission.

Our transcriptional data provide additional evidence that pgm2 deficiency is perceived as a cell-wall stress. Bulk RNA sequencing revealed coordinated induction of stress/carbon-response genes together with components of the cell-wall integrity pathway, including *RHO1*, *SLT2/MPK1*, and *RLM1*, and the remodeling/structural genes *CRH1, GAS1* and *PIR3* (38–40). These responses were broadly supported by qRT-PCR and were largely reversed by either *Pjpgm2* or *Pmpgm2* complementation. This pattern is consistent with activation of the fungal cell wall-integrity (CWI) response rather than an isolated change in glucan synthase expression.

These findings also connect the present study to earlier studies of *Pneumocystis* cell-wall signaling. Thomas and colleagues characterized *P. carinii* Bck1, an upstream component of the fungal CWI MAP kinase cascade and demonstrated that *Pcbck1* complements the cell-lysis phenotype of *S. cerevisiae bck1*Δ cells and restores stress-dependent *FKS2* expression (41, 42). A *Pneumocystis* Mkp1 MAP kinase homologous to the yeast Slt2/Mpk1 cell-integrity kinase has also been characterized (43, 44). Other studies identified environmentally responsive PcCbk1 and demonstrated signaling between PcSte20 and PcCbk1 during interactions with lung epithelial cells and extracellular matrix (45). Collectively, these studies indicate that *Pneumocystis* possesses conserved signaling mechanisms that sense environmental and cell-wall stress and couple these signals to wall remodeling.

The remodeling response observed here also has direct links to other previous *Pneumocystis* studies. Kottom et. al characterized *P. carinii phr1*, a homolog of *S. cerevisiae GAS1* and *Candida PHR1/PHR2* (46). PcPhr1 is pH responsive and belongs to a family of proteins implicated in β-glucan cross-linking and maintenance of cell-wall integrity (47). The reduction of *GAS1* transcription observed in *pgm2*-deficient yeast, followed by restoration towards WT + vector cells with *Pjpgm2* and *Pmpgm2*, therefore links the current transcriptional phenotype to a glucan-remodeling system already demonstrated in *Pneumocystis*. Together with the earlier Gsc1 studies, these findings suggest that PGM-dependent metabolism may influence both production of the β-1,3-glucan backbone and its subsequent organization within the fungal wall.

An important distinction, however, is that *S. cerevisiae* possesses compensatory pathways that are reduced or absent in *Pneumocystis*. Comparative genomic and biochemical studies demonstrated that *Pneumocystis* lacks chitin synthases and chitinases and does not contain detectable chitin (48). β-glucan is therefore retained in a highly reduced cell-wall biosynthetic repertoire lacking several structural polymers used by other fungi. This reduced redundancy may make the maintenance of normal β-glucan levels particularly important in the cyst form and may limit the ability of *Pneumocystis* to compensate when UDP-glucose availability or β-glucan synthesis is disrupted.

A similar distinction should be made for *FKS2/GSC2*. In *S. cerevisiae*, Fks1 and Fks2 provide partially overlapping β-1,3-glucan synthase functions, with *FKS2* serving as an important stress-responsive component of the CWI program (49–51). In the present study, *FKS2/GSC2* was strongly induced by pgm2 deficiency in the RNA-seq dataset. qRT-PCR showed the same directional increase and reversal following complementation, although the qPCR differences did not reach statistical significance. Thus, *FKS2* induction is best interpreted as evidence of a compensatory response to glucan/cell-wall stress rather than as proof that increased Fks2 activity itself accounts for the phenotype. In *Pneumocystis*, only the Gsc1/Fks1 glucan synthase has been functionally characterized, further emphasizing that the exact compensatory architecture identified in yeast should not be assumed to occur identically in *Pneumocystis*.

The caspofungin results provide an important functional test of this model. Despite induction of the stress-responsive *FKS2/GSC2* pathway, *pgm2*Δ yeast was markedly more susceptible to low-dose caspofungin, whereas *Pjpgm2* and *Pmpgm2* complementation restored susceptibility toward WT + vector cells. These observations suggest that transcriptional activation of the Fks pathway is insufficient to compensate when the metabolic support for glucan synthesis is impaired. One interpretation is that glucan synthase abundance and glucan precursor availability represent complementary constraints on wall production, increasing the synthase machinery cannot fully restore wall homeostasis when PGM-dependent carbohydrate generation is compromised.

This concept has particular relevance to emerging echinocandin resistance. Recent work in *Candida glabrata* has directly connected the CWI kinase Slt2 with Fks2-mediated echinocandin resistance; disruption of *SLT2* reduced *FKS2* expression and abolished resistance associated with Fks2, but not Fks1, mutations (50). Even more recent studies indicate that calcineurin, Slt2, Rlm1, and other stress-response regulators contribute to Fks2-associated echinocandin resistance (52). Thus, Fks2 is increasingly understood as part of a broader adaptive network rather than simply an alternative glucan synthase. Our finding that Pgm2 deficiency activates CWI signaling and increases *FKS2* expression, yet renders cells more susceptible to caspofungin, suggests that this compensatory response is insufficient to maintain cell-wall integrity when Pgm2 activity is disrupted.

The relationship between PGM activity and glucan homeostasis is also emerging in other pathogenic fungi. Genetic reduction of PGM in *Aspergillus fumigatus* impairs fungal growth and cell-wall integrity, and recent analysis of a conditional *A. fumigatus pgm* (*Af*PGM) mutant demonstrated decreased soluble β-glucan and impaired biofilm formation (53). The similar reduction in β-glucan observed following *Af*PGM disruption in *A. fumigatus* and Pgm2 deficiency in the present yeast model suggests a conserved role for PGM activity in maintaining normal β-glucan levels across diverse fungal species.

These findings strengthen the rationale for fungal PGM as a therapeutic target. ISFP10 was developed from an isothiazolone scaffold that covalently targets a fungal PGM cysteine absent from the corresponding human enzyme (54). In *A. fumigatus*, the critical residue is C353 (54); importantly, our previous sequence analysis identified the corresponding cysteine at C351 in PjPgm2 and C352 in PmPgm2 (10, 11), whereas this residue is absent from human and mouse Pgm1. ISFP10 inhibits *Af*PGM with an IC_50_ of approximately 2 μM and shows at least 50-fold selectivity relative to human PGM (54). These species-specific residues provide a structural basis for selective pharmacologic targeting of fungal PGM.

The therapeutic evidence has also expanded considerably. In addition to inhibition of *Aspergillus* and *Pneumocystis* Pgm2, recent studies demonstrate ISFP10 activity against *Candida albicans* and *C. glabrata* (30). Short-term administration in mice was generally well tolerated without evidence of major organ toxicity, further supporting continued development of this scaffold (30). More recent *Pneumocystis* studies have also reported inhibition of PjPgm2 and PmPgm2 by ISFP10 and reduction of *P. murina* burden in experimental systems (10), strengthening the translational rationale for targeting this pathway.

The present findings raise an additional therapeutic possibility: PGM inhibition may both directly restrict fungal carbohydrate metabolism and weaken the ability of the cell wall to withstand inhibition of β-1,3-glucan synthesis. This creates a potential two-pronged approach in which metabolic limitation of UDP-glucose supply is combined with inhibition of its downstream utilization by glucan synthase. The enhanced caspofungin susceptibility of *pgm2*Δ yeast provides proof of principle for this concept, although pharmacologic combination studies with ISFP10 and echinocandins will be required to establish synergy. Such studies may be particularly relevant in fungi in which CWI/Fks2 adaptation contributes to echinocandin tolerance or resistance. Echinocandin resistance remains especially important in *C. glabrata* (55), while emerging Fks-dependent resistance and resistance reinforce the need for strategies that disable compensatory wall-stress pathways (56).

For *Pneumocystis*, targeting glucan metabolism may have additional biological consequences because β-1,3-glucan is closely linked to the cyst stage (6). The cyst wall contains abundant β-glucan (6, 57, 58), whereas trophic forms largely lack this polymer, and β-glucan synthase inhibition markedly depletes cysts (35–37). Because the cyst is associated with transmission (59) and because released β-glucans also contribute to host inflammatory responses (3, 60), interference with PGM-dependent glucan metabolism could potentially influence wall integrity, life-cycle progression, transmission, and host-pathogen interactions. These possibilities will require direct testing in *Pneumocystis* models.

The use of *S. cerevisiae* provides a useful model for investigating *Pneumocystis* Pgm2 function because continuous cultivation and genetic manipulation of *Pneumocystis* remain challenging. However, important differences in cell-wall biology should be considered when extending these findings to *Pneumocystis*. Unlike *S. cerevisiae*, *Pneumocystis* lacks chitin and has only one functionally characterized β-1,3-glucan synthase (28), limiting the compensatory pathways available during cell-wall stress. Accordingly, the *FKS2*-mediated response observed in yeast may not occur in the same manner in *Pneumocystis*, and the caspofungin experiments demonstrate increased dependence on β-1,3-glucan synthase activity rather than a specific requirement for Fks2. Nevertheless, the ability of both PjPgm2 and PmPgm2 to restore β-glucan abundance, zymolyase susceptibility, transcriptional responses, and caspofungin tolerance provides complementary biochemical and functional evidence linking these phenotypes to conserved PGM activity.

In conclusion, the present findings extend previous characterization of *Pneumocystis* Pgm2 by placing this enzyme within a broader network connecting central carbohydrate metabolism to fungal cell-wall synthesis and stress adaptation. Loss of Pgm2 reduces β-1,3-glucan, alters wall function, activates CWI and remodeling responses, and increases susceptibility to β-1,3-glucan synthase inhibition. Restoration by both *Pneumocystis* Pgm2 homologs demonstrates functional conservation of this relationship. When considered alongside the established *Pneumocystis* Gsc1 glucan synthase, Phr1/Gas1-like remodeling system, and conserved CWI signaling machinery, these data support a model in which PGM-dependent metabolism plays an important upstream role in maintaining normal β-glucan levels. The presence of selectively targetable fungal PGM residues, expanding activity of ISFP10 across pathogenic fungi, and the potential to increase vulnerability to cell-wall-directed therapy together provide a strong rationale for continued development of fungal PGM as an antifungal target.

## FINANCIAL SUPPORT

This work was supported by the National Institutes of Health [grants R21AI203622-01 and R01HL062150-33 to A.H.L].

## CONFLICT OF INTEREST

The authors declare no conflicts of interest.

## ETHICAL STATEMENT

This study was performed in strict accordance with the recommendations in the Guide for the Care and Use of Laboratory Animals (61). The Mayo Clinic Institutional Animal Care and Use Committee approved this research (A00008661-26).

## DATA AVAILABILIITY

The RNA-sequencing data generated in this study, including raw sequencing reads and processed gene-expression data, are available through the NCBI Gene Expression Omnibus (GEO) and Sequence Read Archive (SRA) under accession number [accession number]. Additional data supporting the findings of this study are provided within the article and its Supplementary Information.

## Supplementary Material for *Pneumocystis* phosphoglucomutase links glucose-phosphate metabolism to β-1,3-glucan synthesis and cell-wall integrity

**Supplementary Table S1.**
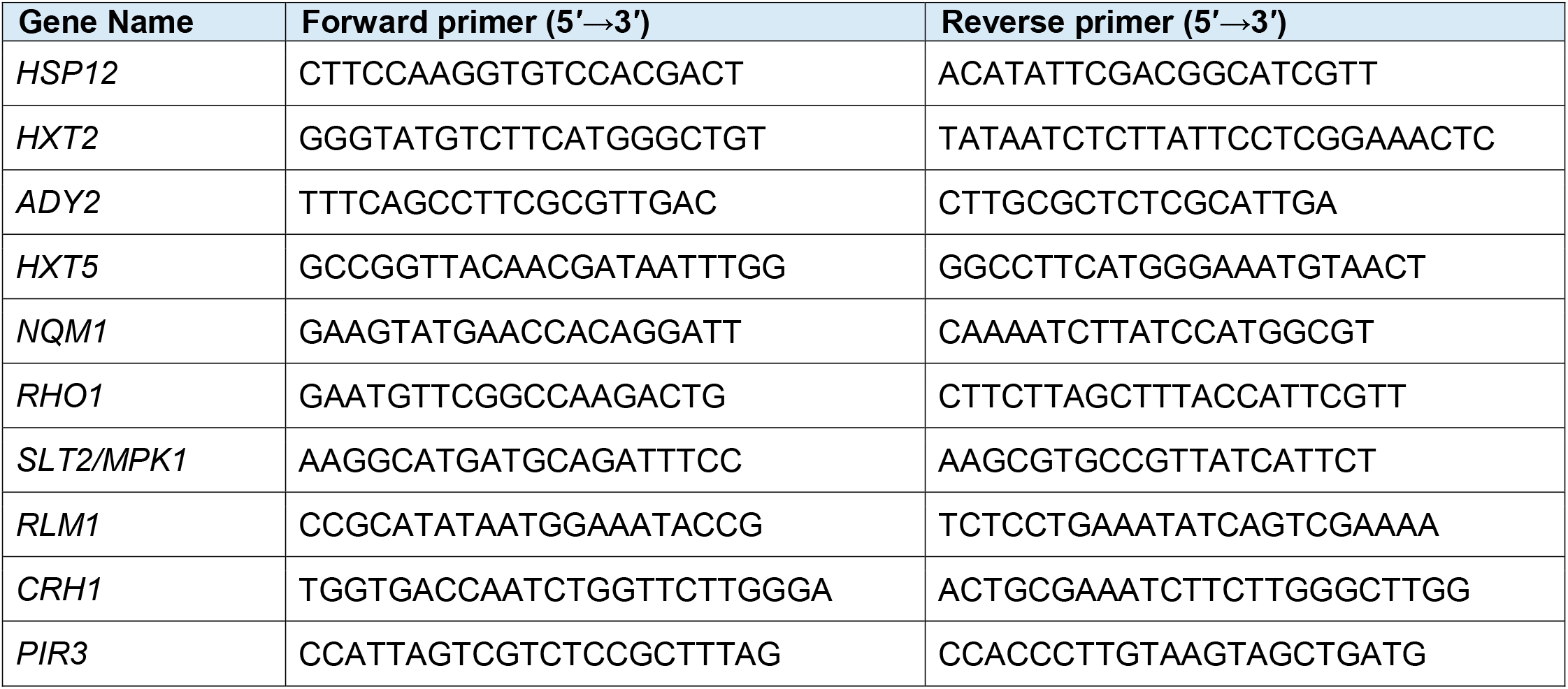
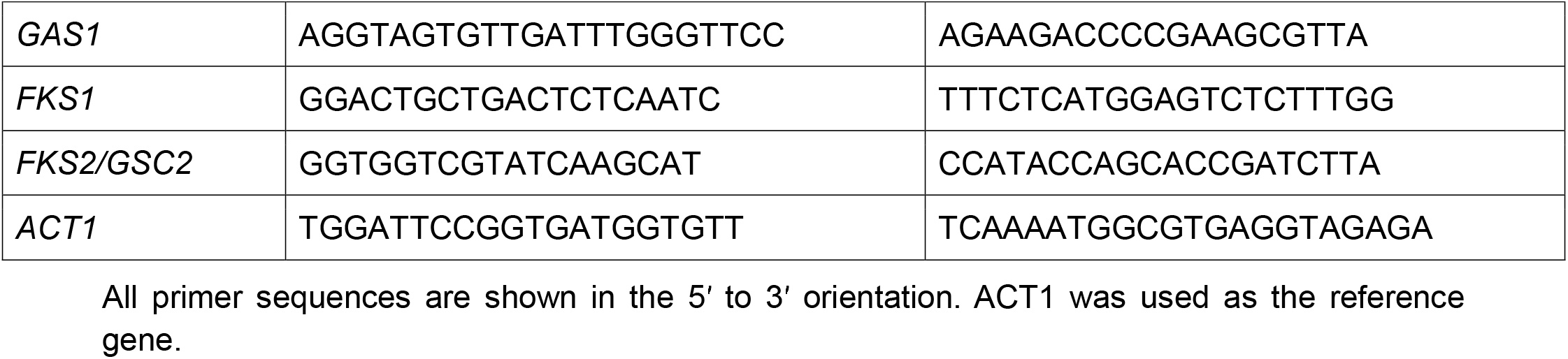
*Saccharomyces cerevisiae* qPCR primer list.

## Notes

### Competing Interest Statement

The authors have declared no competing interest.

